# A single exposure to altered auditory feedback causes observable sensorimotor adaptation in speech

**DOI:** 10.1101/2021.07.26.453857

**Authors:** Lana Hantzsch, Benjamin Parrell, Caroline A. Niziolek

**Author notes:** Equal contribution (Benjamin Parrell, Caroline A. Niziolek).

## Abstract

Sensory errors caused by perturbations to movement-related feedback induce two types of behavioral changes that oppose the perturbation: rapid compensation within a movement, as well as longer-term adaptation of subsequent movements. Although adaptation is hypothesized to occur whenever a sensory error is perceived (including after a single exposure to altered feedback), adaptation of articulatory movements in speech has only been observed after repetitive exposure to auditory perturbations, questioning both current theories of speech sensorimotor adaptation as well as the universality of more general theories of adaptation. Thus, positive evidence for the hypothesized single-exposure or “one-shot” learning would provide critical support for current theories of speech sensorimotor learning and control and align adaptation in speech more closely with other motor domains. We measured one-shot learning in a large dataset in which participants were exposed to intermittent, unpredictable auditory perturbations to their vowel formants (the resonant frequencies of the vocal tract that distinguish between different vowels). On each trial, participants spoke a word out loud while their first formant was shifted up, shifted down, or remained unshifted. We examined whether the perturbation on a given trial affected speech on the subsequent, unperturbed trial. We found that participants adjusted their first formant in the opposite direction of the preceding shift, demonstrating that learning occurs even after a single auditory perturbation as predicted by current theories of sensorimotor adaptation. While adaptation and the preceding compensation responses were correlated, this was largely due to differences across individuals rather than within-participant variation from trial to trial. These findings are more consistent with theories that hypothesize adaptation is driven directly by updates to internal control models than those that suggest adaptation results from incorporation of feedback responses from previous productions.

## Introduction

Auditory feedback plays a major role in both the online execution of speech production and the refinement of feedforward speech motor control, as observed when the auditory feedback participants receive about their own speech is perturbed in real time (Houde & Jordan, 1998; Purcell & Munhall, 2006b; Tourville et al., 2008; Villacorta et al., 2007). Two types of behavior have been the primary focus of auditory perturbation studies in speech, which most typically alter a speaker’s vowel formants (the resonant frequencies of the vocal tract that distinguish vowels). First, when unpredictable vowel formant perturbations are delivered, speakers produce a *compensation* response—an online, within-trial adjustment to oppose the perturbation (Purcell & Munhall, 2006b; Tourville et al., 2008). Second, consistent perturbations of auditory feedback lead to *sensorimotor adaptation*—a learned change in speech behavior that is observable from the onset of speech and which persists even after the perturbation is removed (Houde & Jordan, 1998; Purcell & Munhall, 2006a).

These behaviors are widely considered to be driven by sensory prediction errors (differences between expected and perceived sensory feedback), although models differ in the proposed mechanism by which this occurs. In the DIVA (Directions Into Velocities of Articulators) model (Tourville & Guenther, 2011), sensory prediction errors lead to feedback-based corrective motor commands (i.e. the compensation response) which are subsequently incorporated into the feedforward motor program used for future productions of the same syllables, creating the adaptation response (Kawato et al., 1987). An alternative theoretical account of adaptation (Houde & Nagarajan, 2011) suggests sensory prediction errors instead directly lead to updates of internal models in the sensorimotor control system, either to forward models predicting the sensory outcomes of actions (Bastian, 2006; Haith & Krakauer, 2013; Houde & Nagarajan, 2011; Krakauer & Mazzoni, 2011; Shadmehr et al., 2010), to the control policy guiding action (Hadjiosif et al., 2020), or to both (Wolpert et al., 1998; Wolpert & Kawato, 1998).

Both the compensation-based and internal-model hypotheses of sensorimotor adaptation predict that learning in speech occurs continuously, such that changes in speech production should be evident even after a single trial with altered auditory feedback. Such *one-shot adaptation* has been observed in limb control, where a visuomotor perturbation on an isolated trial affects reach direction on the following trial (Diedrichsen et al., 2005; Joiner et al., 2017; Ruttle et al., 2021). However, the occurrence of such one-shot adaptation has not been definitively established in speech. Although Cai and colleagues (Cai et al., 2012) observed that first formant (F1) production in the first 50 ms of perturbed trials which closely followed another perturbed trial tended to oppose the preceding perturbation’s direction, more recent work explicitly testing for such single trial effects did not find evidence of a measurable change (Daliri et al., 2020). This failure to find one-shot adaptation in speech questions both current theories of speech sensorimotor adaptation as well as the universality of domain-general theories (e.g., Houde & Nagarajan, 2011; Kawato et al., 1987; Hadjiosif et al., 2020).

Here, we aim to further investigate the mechanisms underlying sensorimotor adaptation by measuring one-shot adaptation in speech. To detect this potentially small effect, data from six prior studies (Niziolek et al., 2014; Niziolek & Guenther, 2013; Niziolek & Parrell, 2021; Parrell et al., 2017, 2021) were compiled for this analysis (131 total participants, 18-40 participants per study). In all studies, participants read aloud monosyllabic words while receiving real-time auditory playback of their speech. On a given trial, this feedback was either veridical (*unperturbed trials*) or unpredictably perturbed via an upward or downward shift in F1 (*perturbed trials)* (Figure 1A). Perturbed trials were used to calculate compensation responses, and unperturbed trials which occurred directly after a perturbed trial (*post-perturbation trials)* were used to calculate one-shot adaptation responses (see Figure 1B). We hypothesized that F1 frequency values would be higher for trials that occurred directly after a downward perturbation and lower in trials that occurred directly after an upward perturbation, such that they echo the preceding compensation responses’ F1 values.

**Figure 1:**
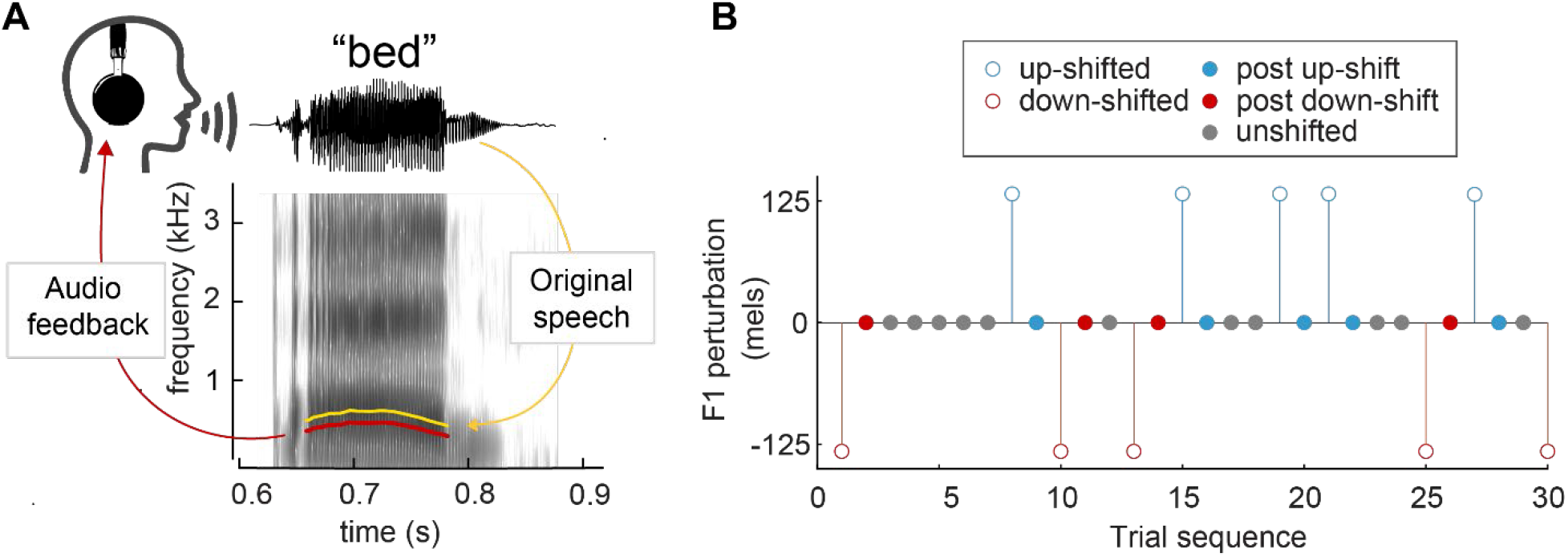
F1 perturbation methodology. **A:** A spectrogram of the word ‘bed’, demonstrating an applied downward F1 perturbation. The F1 frequency of the audio feedback (red) is lowered from the original utterance (yellow). **B:** Sample trial sequence from Study 4. Open circles indicate trials in which an F1 perturbation was applied, and closed circles indicate trials in which no perturbation occurred. “Up-shift” and “down-shift” trials were used to calculate the compensation response. “Post up-shift” and “post down-shift” trials were used to calculate the one-shot adaptation response. For analysis, all shift magnitude values were converted to mels.

This approach also allows us to test the feedback-command-based hypothesis of adaptation in speech, which suggests that there should be a correlation between the magnitude of compensation and subsequent one-shot adaptation at the trial level. While this correlation has been observed in reaching (Albert & Shadmehr, 2016), most studies have failed to identify such a clear relationship in speech (Daliri, 2021; Franken et al., 2019; Lester-Smith et al., 2020; Parrell et al., 2017; Raharjo et al., 2021), possibly because they did not use such a direct trial-to-trial measurement method. The presence of such a relationship at the trial level would be compatible with both the feedback-command-based and internal-model hypothesis of adaptation; alternatively, the absence of such a relationship would only support the internal-model hypothesis.

## Results

### Compensation

In the 150-250 ms time window after vowel onset, perturbed trials in which an upward F1 shift occurred (*up-shifted trials*) had reliably lower F1 values (−3.99 ± 34.13 mels) than trials in which a downward F1 shift occurred (2.69 ± 33.5 mels) (*down-shifted trials*) (*β* = -6.93, S.E. = 0.66, *p* < 0.001, *d* = 0.21). This was also reflected at the individual level; participants’ average F1 in the 150-250 ms time window was substantially lower across up-shifted trials (−2.75 ± 8.68 mels) compared to their average F1 in the same time window across down-shifted trials (4.40 ± 6.96 mels) (paired t-test, *t*(130) = -7.00, *p* < 0.001, *d* = 0.91, Fig 2B, left panel).

**Figure 2:**
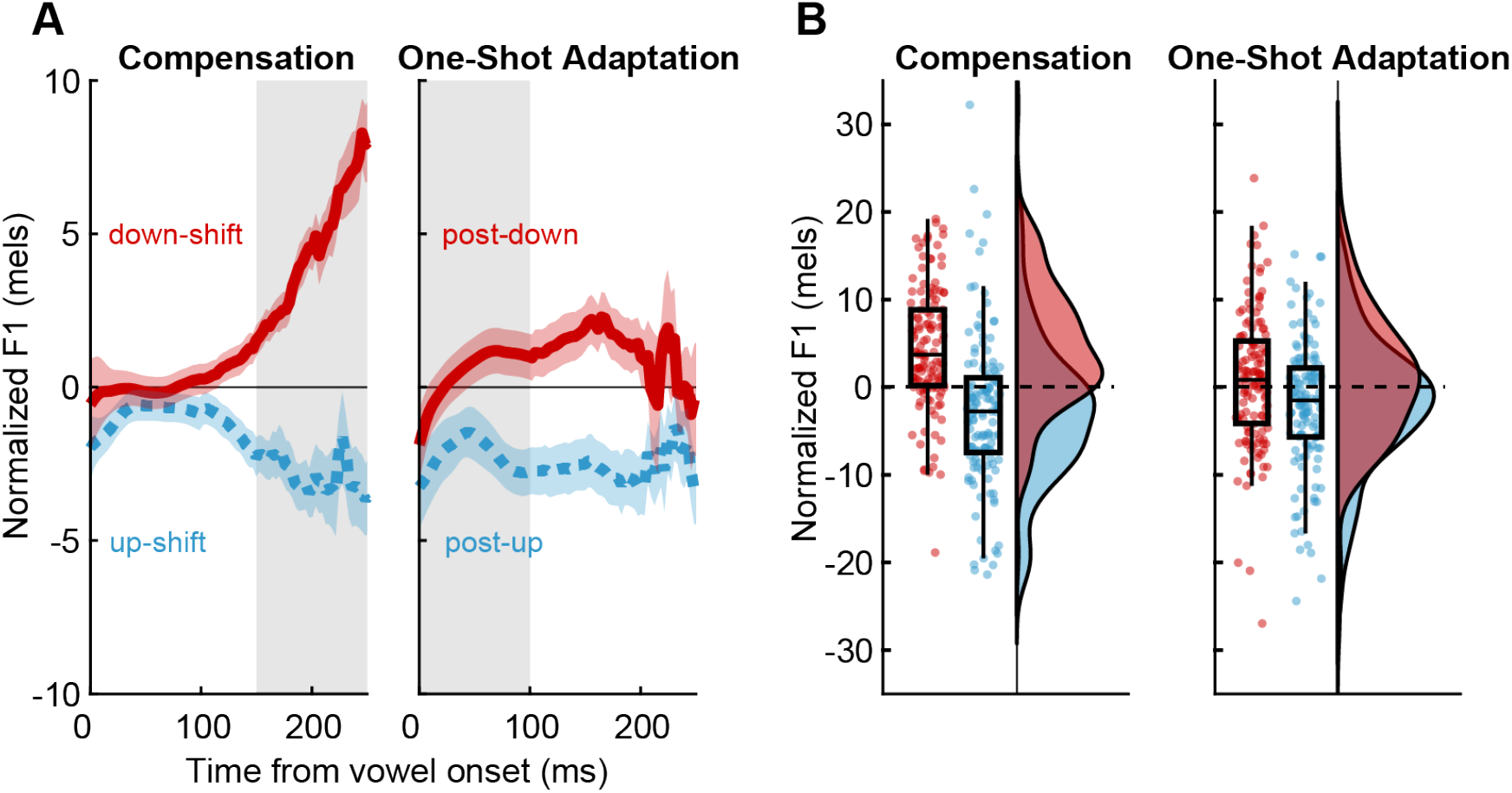
Behavioral responses to auditory perturbations. **A:** The average F1 trajectory for trials that were measured in relation to either upward (blue) or downward (red) perturbations, compiled across participants. Time window of interest is highlighted, illustrating the time period of interest for compensation (left) and one-shot adaptation (right). **B:** The probability distribution and boxplot of participants’ average compensation (left) and one-shot adaptation (right) from trials occurring during or directly after (respectively) up-shifted trials (blue) and down-shifted trials (red).

### One-shot adaptation

Participants produced one-shot adaptation responses which paralleled the directional pattern seen in the compensation response, though at a lower magnitude. In the 0-100 ms time window after vowel onset, F1 values on trials that occurred immediately after an upward F1 shift (−1.55 ± 26.98 mels) were reliably lower than on trials that occurred immediately after a downward F1 shift (0.59 ± 27.8 mels) (*β* = -2.14, S.E. = 0.53, *p* < 0.001, *d* = 0.079). Likewise, participants’ average F1 were lower across trials that occurred directly after an up-shifted trial (−2.08 ± 7.4 mels) than across trials that occurred after a down-shifted trial (0.82 ± 7.73 mels) (paired t-test, *t*(130) = -2.98, *p* = 0.0034, *d* = 0.38, Fig 2B, right panel).

### Relationship between behavioral responses

At the participant level, there was a significant positive relationship between compensation and one-shot adaptation (*β* = 0.14, S.E. = 0.058, *p* = 0.015, η^2^ = 0.02), such that participants who produced larger compensation responses tended to adapt more (Fig. 3A). Conversely, the trial-level model revealed no main effect of compensation response (*β* = - 0.033, S.E. = -0.053, *p* = 0.53) (Fig. 3B). However, we did observe a small but significant interaction between shift magnitude and compensation response (*β* = 0.16, S.E. = 0.052, *p* = 0.0023, η^2^ = 0.0009). Along with the finding that larger shift magnitudes led to larger one-shot adaptation responses (*β* = 7.46, S.E. = 3.34, *p* = 0.03, η^2^= 0.04), this suggests that compensation is predictive of adaptation only at larger shift magnitudes. A post-hoc Monte Carlo simulation confirmed that this effect is unlikely to be caused solely by variation in response magnitude across participants (*p* < 0.05).

**Figure 3:**
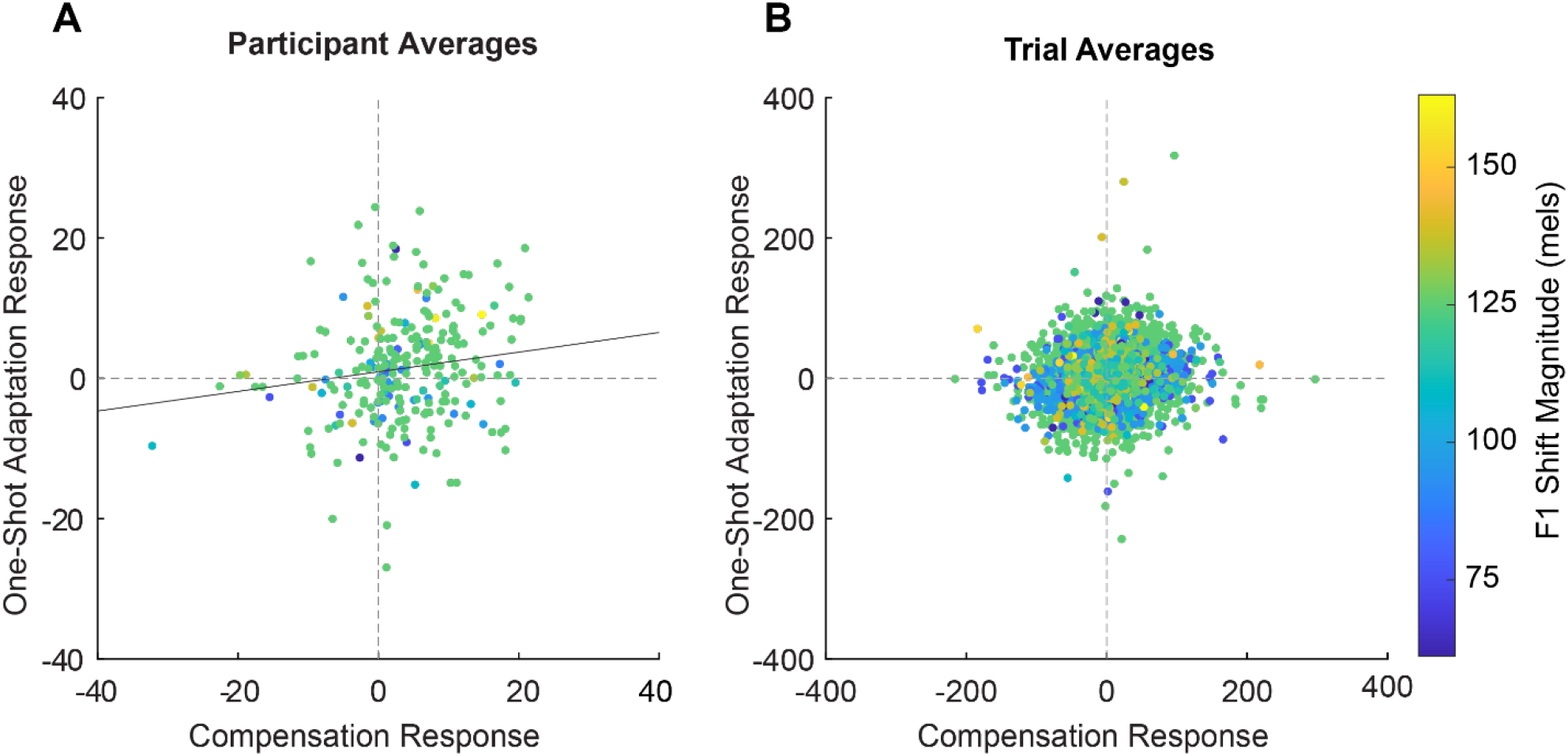
Correlation between compensation and one-shot adaptation. **A:** The correlation between participant average compensation and one-shot adaptation responses. Each participant contributed two data points: their average response to up-shifted and their average response to down-shifted trials. The average applied F1 shift magnitude is displayed via the color gradient (blue = low shift magnitude, yellow = higher shift magnitude). The trend line (y = 0.14x + 0.93) represents the main effect of compensation on one-shot adaptation obtained from the linear mixed model used to analyze this relationship. **B:** The relationship between the trial-level compensation and subsequent one-shot adaptation response. Likewise, the average applied F1 shift magnitude is displayed via the color gradient.

## Discussion

At both the trial and participant level, one-shot adaptation was detected in post-perturbation trials, where F1 values in the first 100 ms of unshifted trials reliably opposed the perturbation in the previous trial. This shows that learning occurs continuously when the sensorimotor system detects a discrepancy between expected and perceived auditory feedback, as predicted by current models of sensorimotor adaptation in speech. While the magnitude of this one-shot adaptation may be small (1-2 mels), it is relatively substantial when accounting for the fact that a typical perturbation of ∼100-150 mels causes a change in F1 an average of only 40-50 mels over the course of 100 or more trials (Katseff et al., 2012; MacDonald et al., 2010; Munhall et al., 2009; Purcell & Munhall, 2006a). Moreover, our estimate of one-shot adaptation is likely conservative. First, most of the studies involved multiple stimulus words in pseudorandom order; in ∼53% of the trial pairs, participants pronounced different words on the perturbed and subsequent unperturbed trial. Although sensorimotor learning can generalize across words with the same vowel (Rochet-Capellan et al., 2012), such generalization is only partial, and a larger adaptation effect likely would have emerged with uniform word pairs. Second, in our planned analysis of one-shot adaptation responses, we measured F1 frequency during the first 100 ms of each vowel in relevant utterances. However, using the 50-150 ms window would have avoided the inclusion of the consonant transition in our measurement. In these data, this would have yielded an average adaptation effect of 2.21 ± 7.81 mels, somewhat larger than that of the values obtained by averaging across the first 100 ms (1.45 ± 7.57 mels).

Though the magnitude of an individual’s average compensation response was predictive of their average one-shot adaptation response, such a general relationship was not reliable at the trial level, where compensation only displayed predictive power at higher shift magnitudes. While theories of adaptation based on changes to internal models would not require the presence of this trial-level relationship, the compensation-based adaptation framework of the DIVA model would predict a larger and more consistent effect. In sum, our results question whether these two behavioral responses have such a direct feedforward relationship, or if this relatively weak correlation could best be explained by compensation and one-shot adaptation responses occurring via separate mechanisms driven by the same sensory error (as may be predicted by internal-model hypotheses).

Overall, these results provide evidence that a single exposure to altered auditory feedback induces “one-shot” adaptation in the speech sensorimotor system. This is consistent with current models of adaptation in speech specifically and in human movement more broadly; within these frameworks, one-shot adaptation is an effect that may continually build upon itself to create more enduring adaptation responses. The expected relationship between compensation and adaptation was observed mainly at the participant, rather than trial, level. While not conclusive, these results are more consistent with models of adaptation that rely on updates to internal models compared to models that use feedback corrections to update future feedforward commands. Our results provide evidence that adaptation in speech may operate in a similar manner as in other motor domains. As a well-learned natural behavior that relies primarily on implicit learning, speech offers a unique, ecologically valid paradigm to further our understanding of the underlying mechanisms driving sensorimotor adaptation.

## Acknowledgements

This work was funded by National Institutes of Health grants R01 DC017091 (B.P.) and R00 DC014520 (C.A.N.).

## Methods

### Participants

We reanalyzed data from six previous studies examining online compensation responses to formant frequency alterations with similar speech stimuli and perturbation schedules. Data were included if participants met inclusion criteria for their respective studies and if the formant shifts they received were opposite or near-opposite each other (separated by an angle of 180 ± 20° when plotted together in F1/F2 space). Data from 91 participants met these criteria; 40 of these participants contributed to two of the included studies. All participants were native speakers of American English and reported no history of speech, hearing, or neurological disorders.

### Auditory perturbation

Details of the six studies are provided in Table 1. In all studies, participants spoke aloud monosyllabic English words containing the vowel /ε/ (as in *head*), which were presented as text on a screen. Simultaneously, participants heard real-time auditory feedback of their speech through headphones. On a pseudorandom subset of trials (25-50%), auditory feedback was altered with one of two real-time feedback perturbation systems, Audapter (Cai et al., 2008; Tourville et al., 2013) or Feedback Utility for Speech Production (FUSP) (Katseff et al., 2012; Parrell et al., 2017) (Fig. 1). Briefly, linear predictive coding (LPC) was used to model the vowel portion of the signal and apply a formant shift in real time during speech. Unaltered trials (50-75% of trials) underwent the same processing pipeline but with no alteration to the formants, such that auditory feedback in all trials had the same (minimal) delay. The magnitude and direction of the applied formant shift varied slightly across studies. Studies 1, 2, 3, and 4 shifted F1 upward and downwards at a consistent magnitude (in mels or Hz) that was applied to all participants. Studies 5 and 6 each calculated participant-specific shift magnitudes for both F1 and F2 (in mels or Hz) along a vector pointing from the target vowel /ε/ to adjacent vowels /I/ (as in *hid*) and /æ/ (as in *had*). For these studies, only the F1 portion of the vector was considered in the analysis; perturbations that increased F1 (/ε/ to /æ/) were considered “up” shifts and perturbations that decreased F1 (/ε/ to /I/) were considered “down” shifts. All formant values were converted into mels for purposes of this analysis.

**Table 1.**
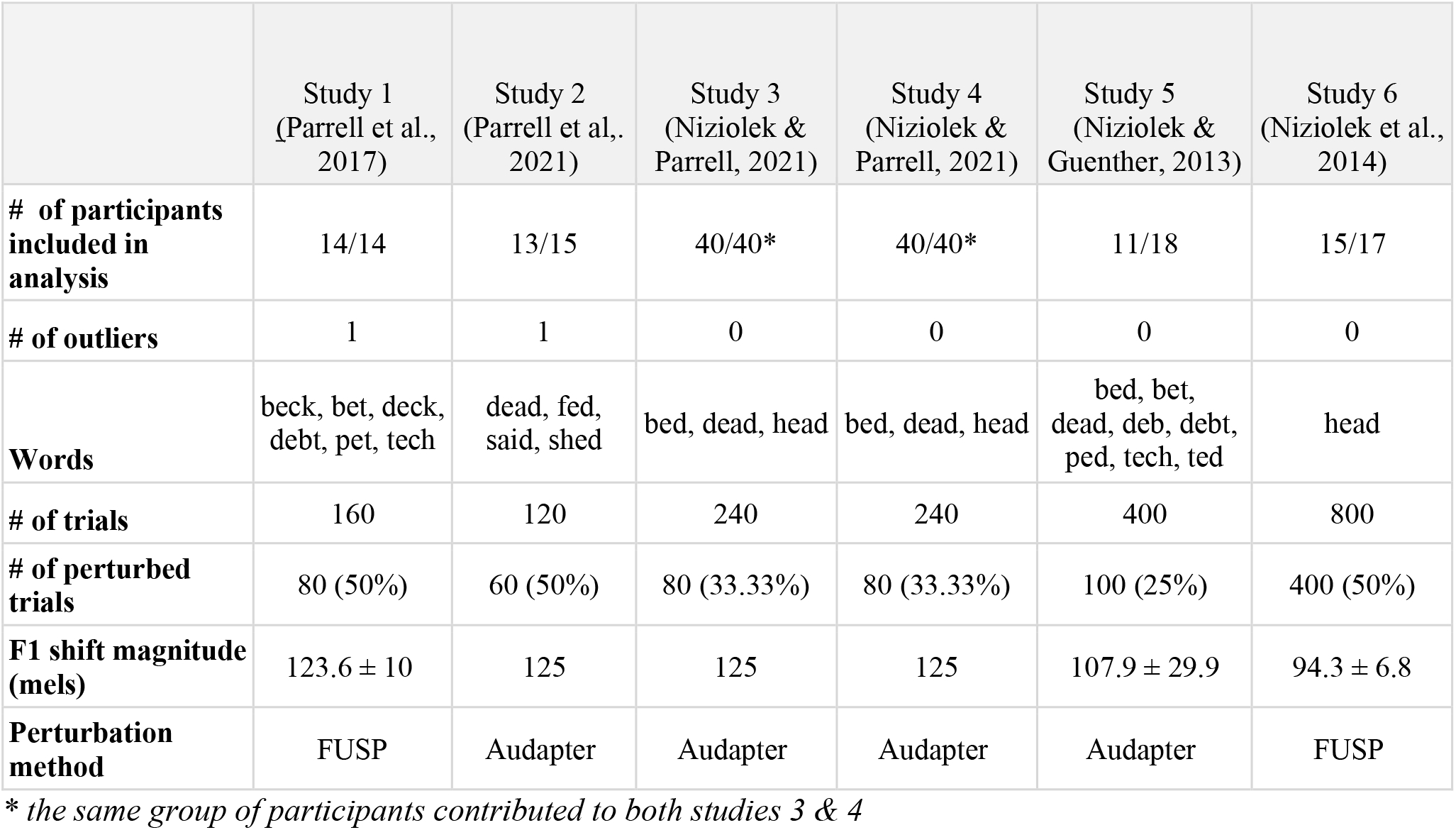
Summary of the included studies.

|  | Study 1<br>(Parrell et al.,<br>2017) | Study 2<br>(Parrell et al.,<br>2021) | Study 3<br>(Niziolek &<br>Parrell, 2021) | Study 4<br>(Niziolek &<br>Parrell, 2021) | Study 5<br>(Niziolek &<br>Guenther, 2013) | Study 6<br>(Niziolek et al.,<br>2014) |
| --- | --- | --- | --- | --- | --- | --- |
| **# of participants**<br>**included in**<br>**analysis** | 14/14 | 13/15 | 40/40* | 40/40* | 11/18 | 15/17 |
| **# of outliers** | 1 | 1 | 0 | 0 | 0 | 0 |
| **Words** | beck, bet, deck,<br>debt, pet, tech | dead, fed,<br>said, shed | bed, dead, head | bed, dead, head | bed, bet,<br>dead, deb, debt,<br>ped, tech, ted | head |
| **# of trials** | 160 | 120 | 240 | 240 | 400 | 800 |
| **# of perturbed**<br>**trials** | 80 (50%) | 60 (50%) | 80 (33.33%) | 80 (33.33%) | 100 (25%) | 400 (50%) |
| **F1 shift magnitude**<br>**(mels)** | 123.6 ± 10 | 125 | 125 | 125 | 107.9 ± 29.9 | 94.3 ± 6.8 |
| **Perturbation**<br>**method** | FUSP | Audapter | Audapter | Audapter | Audapter | FUSP |

\* the same group of participants contributed to both studies 3 & 4

### Behavioral measures and statistical analysis

Our primary measure of interest was one-shot adaptation, an adaptive response that persists in the trial following an isolated perturbation. In order to examine whether one-shot adaptation is related to feedback-based corrections on the previous trial, we additionally measured the online compensation response. These behavioral responses were examined at both the trial level and the participant level.

Trials with a length of less than 100 ms were excluded from analysis (<1%). Two participants were excluded from the analysis as outliers (average compensation or one-shot response > 4 S.D. from mean).

### Compensation

At the *trial level*, compensation response was operationalized as the mean normalized F1 produced during the 150-250 ms time window of trials in which a perturbation occurred (*perturbation trials*). More specifically, participant- and word-specific baseline F1 trajectories were first calculated from the F1 trajectories of unperturbed trials (*baseline trials*). The F1 trajectory of each perturbation trial was then normalized by subtracting the word-specific baseline mean F1 trajectory from it. The compensation response for each perturbation trial was then defined as the mean F1 value within 150-250 ms after vowel onset, after the typical onset latency of compensation. A 200-300 ms time window was originally planned for this analysis; however, only 46% of produced vowels had a duration of at least 300 ms, whereas 80% of vowels lasted until the end of the 150-250 ms time window.

Average compensation response was also calculated at the *participant level*, operationalized as a participant’s mean normalized F1 across the 150-250 ms window of their perturbation trials. Again, the F1 trajectory of each perturbation trial was normalized via a participant- and word-specific baseline. Then for each participant, two average F1 trajectories were calculated: one trajectory that averaged the normalized trajectories across all trials containing an upward perturbation and one trajectory that averaged across all trials containing a downward perturbation. The participant’s average compensation response for each perturbation direction (up and down) was calculated as the mean F1 value in the 150-250 ms time window after vowel onset of these averaged perturbation trajectories.

In the *trial level* analysis, a linear mixed model was employed to investigate the effect of perturbation direction on compensation response: *Compensation response ∼ perturbation direction + (1* | *participant) + (1* | *study)*. Effect size was calculated by dividing β by the residual standard deviation. At the *participant level*, a paired t-test was used to evaluate the distribution of participants’ mean compensation response to upward perturbations vs. downward perturbations. Cohen’s D was calculated to determine effect size.

### One-shot adaptation

At the *trial level*, one-shot adaptation response was calculated as the mean normalized F1 produced in the first 100 ms of unperturbed trials that occurred directly after a perturbed trial (*post-perturbation trials*). Again, participant- and word-specific baseline trajectories were calculated, though using F1 trajectories from unperturbed trials that directly followed another unperturbed trial (*baseline trials*). The F1 trajectories of each post-perturbation trial were then normalized by subtracting the word-specific baseline mean F1 trajectory. The one-shot adaptation response for each post-perturbation trial was calculated as the mean F1 value in the first 100 ms of the normalized trajectory. Only F1 values from the initial 100 ms of the vowel were included, limiting the influence of auditory-based feedback control mechanisms, which have a latency of 100-150 ms in speech (Cai et al., 2012; Parrell et al., 2017; Tourville et al., 2008).

At the *participant level*, the one-shot adaptation response was calculated as a participant’s mean normalized F1 in the first 100 ms of their average post-perturbation trial F1 trajectory. Again, the F1 trajectory of each post-perturbation trial was normalized via a participant- and word-specific baseline. Then for each participant, two average F1 trajectories were calculated: one trajectory that averaged the normalized trajectories across all trials that occurred after an upward perturbation and one trajectory that averaged across all trials that occurred after a downward perturbation. The participant’s average one-shot adaptation response for each perturbation direction (up and down) was calculated as the mean F1 value in the first 100 ms of these averaged post-perturbation trajectories.

At the *trial level*, a linear mixed model was employed to investigate the effect of perturbation direction on one-shot adaptation: *One-shot adaptation response ∼ perturbation direction + (1* | *participant) + (1* | *study)*. Effect size was calculated by dividing β by the residual standard deviation. At the *participant level*, a paired t-test was implemented to assess the distribution of participants’ mean one-shot adaptation response to upward perturbations vs. downward perturbations. Cohen’s D was calculated to determine effect size and conduct a power estimation.

### Relationship between behavioral responses

In order to assess the relationship between compensation and the one-shot adaptation that followed it, we fitted a linear mixed-effects model to one-shot adaptation with compensation, perturbation magnitude, and perturbation condition as fixed factors and with participant as a random intercept. Separate analyses were conducted at the participant level (averaging across all trials) and at the individual trial level. To avoid problems in the linear models caused by predictors of very different scales, each perturbation magnitude was normalized by dividing by the mean of all perturbation magnitudes across participants. Study was not included as a separate random intercept in the model as it introduced singularity to the model due to its collinearity with participant and shift magnitude. In order to remove the directional difference between up and down perturbation conditions and maintain standardized magnitude measure between the two perturbation directions, compensation and one-shot adaptation responses from upward-shifted trials were multiplied by -1.

At the trial level, compensation response was intended to be included as a random slope by participant, however was removed because the model failed to converge. In order to determine whether this trial level relationship was just a reflection of the participant level distribution and not specific to the trial-to-trial relationships, a Monte Carlo simulation was run on the trial level model. For each participant, a random sample of one-shot adaptation and compensation responses was taken from a set of normal distributions. These distributions were calculated based on that participant’s mean and standard deviation of responses separately in the respective up and down shifted conditions. The extracted random samples were then run through the same statistical tests as the original trial level dataset. This simulation was run 1000 times. The resulting distribution of η^2^ values revealed that an effect size of the magnitude observed in the original dataset (η^2^ = 0.0009) occurred in <1% of the random samples (95% = 0.000427).

All statistical analysis was conducted in R (R Core Team, 2020). Linear mixed effects models and their simplest explanatory models (calculated via stepwise regression) were generated using the *lme4* package (Bates et al., 2015). Statistical significance of the final model was assessed with the *lmerTest* package, which uses the Satterthwaite method to estimate degrees of freedom (Kuznetsova et al., 2017). Power analyses for t-tests were conducted with the *pwr* package (Champely, 2020). Correlation between compensation and one-shot adaptation was then assessed with a Pearson R correlation coefficient using the *MuMIn* package (Barton, 2020). Effect sizes were calculated using the *effectsize* package (Ben-Shachar et al., 2020). Data and analysis code is available at https://github.com/blab-lab/postMan. Some of the functions rely on additional code available at https://github.com/carrien/free-speech.

